# PrimPol is required for the maintenance of efficient nuclear and mitochondrial DNA replication in human cells

**DOI:** 10.1101/501304

**Authors:** Laura J. Bailey, Julie Bianchi, Aidan J. Doherty

## Abstract

Eukaryotic Primase-Polymerase (PrimPol) is an enzyme that maintains efficient DNA duplication by repriming replication restart downstream of replicase stalling lesions and structures. To elucidate the cellular requirements for PrimPol in human cells, we generated PrimPol-deleted cell lines and show that it plays key roles in maintaining active replication in both the nucleus and mitochondrion, even in the absence of exogenous damage. Human cells lacking PrimPol exhibit delayed recovery after UV-C damage and increased mutation rates, micronuclei and sister chromatin exchanges but are not sensitive to genotoxins. PrimPol is also required during mitochondrial replication, with PrimPol-deficient cells having increased mtDNA copy number but displaying a significant decrease in replication rates. Deletion of PrimPol in XPV cells, lacking functional Pol Eta, causes an increase in DNA damage sensitivity and pronounced fork stalling after UV-C treatment. We show that, unlike canonical TLS polymerases, PrimPol is important for allowing active replication to proceed, even in the absence of exogenous damage, thus preventing the accumulation of excessive fork stalling and genetic mutations. Together, these findings highlight the importance of PrimPol for maintaining efficient DNA replication in unperturbed cells and its complementary roles, with Pol Eta, in damage tolerance in human cells.

## Introduction

To successfully maintain genome integrity, cells must accurately and efficiently replicate their DNA to pass on accurate copies to daughter cells. During this process, they must deal with lesions that arise due to replication errors or DNA damaging agents, as well as DNA / RNA structures present in the genome. To overcome these obstacles, cells possess a wide range of tolerance and repair pathways, as well as checkpoints, that limit damaged DNA being passed on to daughter cells. Lesions are repaired by several different pathways including nucleotide and base excision repair to remove lesions, mismatch repair to excise incorrectly matched base pairs and HR / NHEJ to repair double-strand breaks (DSBs) (reviewed in (1)). However, when the replication machinery encounters damaged or structured DNA, it must overcome these obstacles to avoid generating breaks, which may lead to the loss of genetic information. To achieve this, cells employ a number of damage tolerance DNA polymerases that can replicate across a range of different lesions in a process termed <u>T</u>ranslesion <u>S</u>ynthesis or TLS (2,3). These included the Y family polymerases Pol Eta, Kappa, Iota, Zeta and Rev1 (2,4). These enzymes have specialised roles in bypassing a range of lesions (4-6). For example, Pol Eta can bypass UV induced cyclopyrimidine dimers (CPDs) and loss or mutation of this gene causes Xeroderma Pigmentosum (XP), a disease characterised by UV sensitivity (7,8). Others, such as Pol Theta, can instigate micro-homology mediated end-joining in order to rejoin and fill in DSBs (9).

Recently, an additional damage tolerance replicase has been identified called Primase-Polymerase, (PrimPol), a member of the archaeal eukaryotic primase (AEP) family (10-14). PrimPol possesses both primase and polymerase activities and is able to bypass a variety of lesions and structures, as well as repriming replication restart at sites of stalled synthesis (10,15-18). A number of studies have shown that PrimPol is important for the maintenance of replication after damage and loss of the protein causes UV-C sensitivity, slowing of replication and cell cycle arrest in avian DT40 cells (10,12,15,19). PrimPol has been shown to interact with a number of replication-associated proteins, such as RPA and PolDIP2, which are likely to be important for its recruitment and function at sites of stalled replication (20-22).

As well as nuclear DNA maintenance, PrimPol is also found in mitochondria (mt) where it is involved in the replication of mtDNA (11,23,24). Unlike in the nucleus, human mitochondria contain multiple copies of a 16 kb circular DNA molecule organised into nucleoids that encodes 13 components of the electron transport chain, 22 tRNAs and 2 rRNAs. mtDNA is replicated by a dedicated polymerase, Pol γ, and a range of other repair and replication proteins are utilised, some of which are mitochondrial-specific and others have dual roles in both the nucleus and mitochondrion (23,25-28). PrimPol has been reported to be important for repriming of mtDNA replication after damage (24), but little is still known about its recruitment to the organelle or its DNA, although it has been shown to interact with mtSSB in a manner functionally similar to RPA (29). Thus, PrimPol is a dynamic protein that likely fulfils similar roles in DNA replication in both the nuclear and mitochondrial compartments.

Here, we extend our understanding of the roles of PrimPol in the maintenance of DNA replication, in both the nucleus and the mitochondrion, with the characterisation of several human cell lines lacking functional PrimPol. We show that PrimPol^-/-^ cells exhibit a small increase in micronuclei, fork stalling, a delay in cell cycle recovery and an increase and change in mutation frequencies after UV-C damage. However, PrimPol^-/-^ cells also show defects in the absence of damage with a decrease in replication fork speeds, micronuclei and sister chromatid exchanges, compared with WT cells. PrimPol-deficient cells also display a significant decrease in mtDNA replication and an increase mtDNA copy number. Although loss of human PrimPol does not overtly affect cell viability after DNA damage, the additional loss of Pol Eta (η) results in increased damage sensitivity, delayed recovery and enhanced fork stalling. Together, these findings highlight the requirement for PrimPol in the maintenance of efficient nuclear and mitochondrial replication in both unperturbed and perturbed human cells.

## Material and Methods

### Cell growth and transfection

MRC5 and XP30RO cells were grown in MEM, 15 % fetal calf serum, 1 % L-glutamine and 1 % penicillin/streptomycin. 143B osteosarcoma cells were grown in DMEM, 10 % fetal calf serum, 1 % L-glutamine and 1 % penicillin/streptomycin, ρ^0^ cells were supplemented with 50 µg/ml uridine. PrimPol^-/-^ and XPV PP1 cells were complemented with PrimPol cloned into pEGFP-C2, with or without the addition of T2a puromycin producing an N-terminally tagged GFP protein. Briefly, cells were transfected with the desired plasmid using lipofectamaine 2000 (Invitrogen) in OPTI-MEM for 4 hrs, before the addition of normal media. After 24 hrs, cells which had taken up the plasmid were selected with G418 (2 mg/ml) or Puromycin (2 µg/ml). Selected populations were then maintained in the selective antibiotic to ensure maintenance of the desired protein. To deplete endogenous PrimPol cells were treated with PrimPol specific or scrambled siRNAs as described previously (10). Cell growth rate was calculated by counting cells at approximately 24 hr intervals using a haemocytometer, growth curves were then used to calculate the doubling time (using Doubling Time Computing, see: http://www.doubling-time.com/compute.php). To monitor cell growth rate in real time cells were analysed in an IncuCyte^™^ phase contrast microscope (Essen Biosciences). After transfection, or at time 0, cells were placed in the microscope and analysed concurrently with 9 regions analysed per sample at 3 hourly timepoints. The IncuCyte 2011A software package was then used to analyse cell density in each image allowing the generation of an average growth curve.

### Disrupting PrimPol using a Zinc Finger gene targeting approach

PrimPol^-/-^ cells were generated using a pre-designed ZFN nuclease (SIGMA) following manufacturer’s instructions. Briefly, MRC5 or XP30RO cells were transfected with mRNA for the PrimPol ZFN using lipofectamine 2000. Activity of the ZFN was confirmed on the pool population compared to mock treated cells using the Surveyor Nuclease assay (Transgenomic) on a PCR product using the supplied primers. Cells were serial diluted and colonies were grown in 96 well plates from single cells. Colonies were screened by PCR with the primers supplied and colonies where products showed size differences were amplified and further screened by western blot using a PrimPol-specific polyclonal antibody raised against purified PrimPol (Eurogentec). Colonies with no observed protein expression were screened further by PCR and a small region across the site of the ZFN nuclease was amplified and separate bands for each clone were gel extracted and sequenced to elucidate changes in the genomic DNA. To confirm deletions, RNA was extracted from cells using Trizol and resuspended in H_2_O. cDNA was generated using M-MuLV reverse transcriptase (NEB) and oligo dT, PrimPol cDNA was amplified by PCR (Phusion NEB) using specific primers (PPfwd, PPrev).

### Flow Cytometry

Cells were analysed by flow cytometry to analyse changes in cell cycle populations. Cells were plated and allowed to attach for at least 16 hrs before being treated with / without UV-C and allowed to recover for the desired time. To mark replicating cells, 10 µM EdU was added to culture media for 25 minutes before cells were collected and fixed with 70 % ethanol. EdU positive cells were labelled using click chemistry with a sulfo-CY5 azide (Jena Biosciences) (30,31). Cells were then treated with 150 µg/ml RNAse A and total DNA was labelled with 5 µg/ml propidium iodide before being separated using a BD Accuri C6 flow cytometer. Approximately 10,000 cells were analysed per sample and cell populations were quantified using the BD CSampler Software. Where initial cell cycle populations were measured then EdU labelling not was used.

### Cell Survival assays

To measure damage sensitivity, 100 cells (or a serial expansion where sensitivity was expected) were plated in standard media and allowed to attach overnight. Increasing concentrations of selected drugs were then added directly to the media or cells were treated with UV-C using a G6T5 Germicidal 9” 6W T5 UVC lamp (General Lamps Ltd.) and allowed to form colonies over ∼10 days. In the case of cisplatin, the drug was washed off 5 hrs after treatment and cells were grown in standard media. 0.38 mM caffeine was added to the media 2 hrs prior to damage, where required. Cells were fixed with ethanol, stained with 1 % methylene blue and counted, with sensitivities based on cell plating efficiencies in undamaged controls.

### HPRT Mutation Assay

The HPRT assay was carried out based on previous methods (32). Approximately 50 × 10^4^ cells were treated with 40 ng/ml 4NQO for 5 days or 10 J/m^2^ UV-C and cells were allowed to recover for 6-7 days. 2 × 10^5^ treated cells were then plated into 96 well plates in the presence of 5 µg/ml 6-thioguanine, or 100 cells with no drug. After 10 days, mutation rates were calculated as the number of colonies formed in the presence of 6-thioguanine based on plating efficiency calculated from cells plated in the absence of the drug. Colonies were then expanded to allow for mutation analysis. RNA was extracted from resistant clones using Trizol and *HPRT* cDNA was generated using M-MuLV reverse transcriptase (NEB) and primer HR1 (as described (33)). The *HPRT* cDNA was amplified using nested PCR with primers HR1 and HF1, followed by HF2 and HR2. PCR products were sequenced by Sanger sequencing with primer HR2 and compared to the reference genotype and control samples from undamaged cells.

### mtDNA copy number and Replication

mtDNA copy number was assessed as described previously (34). Briefly, total cellular DNA was extracted from cells and proteinase K treated. PCR was carried out using a Roche LightCycler 480 and Maxima Probe qPCR reagents (ThermoFisher Scientific) following manufacturers instructions. mtDNA was amplified using a PCR specific to the *CoxII* gene, see Table 1 for primers and 5’ Fam, 3’ Tamra labelled probe. This was quantified by direct comparison with amplification of nuclear DNA with primers to *APP* and a 5’ Fam, 3’ Tamra labelled probe. For DT40 cells, primers and probes were targeted to the nuclear *APP* gene and *CoxII* in mtDNA (see Table 1). mtDNA replication was analysed using BrdU. Nuclear DNA replication was first blocked by the addition of 50 µM aphidicolin for at least 30 minutes and this was followed by the further addition of 10 µM BrdU for a further 24 hrs. DNA was then collected as described above and 200 ng DNA was spotted onto nitrocellulose membrane in triplicate. Once dried, the spots were fixed to the membrane using 0.4 M NaOH. The membrane was then probed with mouse anti-BrdU Clone B44 (BD 347580) and HRP conjugated anti mouse (ab97046) antibodies and results were visualised using ECL (GE Healthcare) and quantified using Image J. To analyse total mtDNA content, the membrane was further probed with a mtDNA specific radiolabelled probe as described below.

### 2D-AGE Analysis

Neutral 2D gel electrophoresis was carried out as described previously (35). Briefly, 30 µg total cellular DNA was digested with HinCII (NEB) and ethanol precipitated. DNA was separated on a 0.4 % agarose gel at 1.1 V/cm for 18 hrs. The lane was then excised from the gel and rotated 90° before being separated in a 1 % agarose gel with 500 ng/ml ethidium bromide for 6 hrs at 260 mA at 4°C. The gel was southern blotted and DNA was visualised using a ^32^P dCTP radio-labelled probe generated using Ready-To-Go™ dCTP DNA labelling beads (GE healthcare) and a PCR product to mtDNA region 16341-151 generated with primers Hs mt16341 and Hs mt151 (see Table 1). The membrane was exposed to a phospho screen which was scanned using a Fujifilm FLA-5100 image reader and images were quantified using AIDA and Image J software.

### Mitotracker analysis

For analysis of mitochondrial membrane potential, cells were stained by the addition of 2.5 µM Mitotracker Green FM or 2.5 µM Mitotracker Red CMXRos (Invitrogen) for 20 mins in MEM only media. Cells were collected and washed with PBS before being analysed on a BD Accuri C6 flow cytometer. For microscopy imaging cells were stained as above with Mitotracker Deep Red before being fixed in 3 % paraformaldehyde and used for immuno-fluorescence. To analyse Mitotracker-stained mtDNA nucleoids, fixed cells were stained with anti-DNA antibody (Progen), followed by anti-mouse Alexa Fluor 488 (Molecular Probes).

### Microscopy imaging

For microscopic imaging, cells were first fixed with 3 % paraformaldehyde. Cells were stained with the relevant antibodies and mounted in ProLong^™^ gold antifade mountant with DAPI (Molecular Probes). Slides were viewed on an Olympus IX70 fluorescent microscope or Nikon E400 fluorescent microscope, images collected using Micro-manager and viewed and quantified with Image J and OMERO.

### Sister Chromatid exchange analysis

SCEs were analysed as described previously (36). Briefly, cells were grown for 48 hrs with the addition of 10 µM BrdU. To increase the number of mitotic cells, 1 µM nocodazole was added for the last 16 hrs before the cells were collected. Cells were swollen in 75 mM KCl at 37°C before being fixed with 3:1 methanol: acetic acid and spotted onto glass slides. After being allowed to dry cells were stained with 10 µg/ml Hoechst. Slides were washed in SSC, (150 mM Sodium Chloride, 15mM Sodium Citrate) and exposed to UV light for 1 hr before being incubated in SSC buffer for a further 1 hr at 60 °C. Finally, slides were stained with Giemsa and after drying mounted with Eukitt Quick-hardening mounting medium (Sigma-Aldrich). Slides were then imaged and counted as described above.

### Fibre analysis

Replication progression was analysed on DNA fibres as described previously (10). Briefly, approximately 15 x10^4^ cells were labelled with 25 µM CldU for 20 minutes followed by 250 µM IdU for a further 20 minutes. For fork stalling assays, a pulse of 20 J/m^2^ UV-C was given in between the two labels. Cells were lysed and DNA was spread down slides using gravity before being fixed with 3:1 methanol: acetic acid. After rehydration, fibres were stained with antibodies to the specific labels, rat anti-BrdU [BU1/75 (ICR1)] (Abcam ab6326), mouse anti-BrdU Clone B44 (BD 347580), anti-rat Alexa Fluor 488 (A21208), anti-mouse Alexa Fluor 594 (A31624) (Invitrogen Molecular Probes). Slides were mounted with Fluoromount (Sigma-Aldrich) and imaged and quantified as described above.

## Results and Discussion

### Generation of a human PrimPol Knockout cell line

Previous human PrimPol depletion studies, using siRNA-knockdown, suggested that it has a role in DNA damage tolerance within the nucleus, as well being involved in the maintenance of mtDNA (10-12,15). Notably, we observed a consistent decrease in cell growth rate after reduction of PrimPol by siRNA, compared to cells treated with a scrambled siRNA control, suggesting an important role in maintaining cell proliferation (Fig. 1A, S1A). As PrimPol has both nuclear and mitochondrial roles, we examined the potential causes of this growth impediment using ρ^0^ cells, which lack mtDNA. As well as having similar levels of endogenous PrimPol (Fig. S1B), ρ^0^ cells exhibited an almost identical decrease in cell proliferation after siRNA depletion of PrimPol, compared to cells treated with a scrambled siRNA control (Fig. 1A, S1C-E). Thus, it appears that loss of PrimPol impacts on cell cycle progress, likely through nuclear DNA maintenance and / or replication. However, PrimPol mice are viable and display no overt phenotypes thus cells must be able to adapt to this loss, ultimately allowing a sufficient proliferation rate to be maintained following loss of PrimPol (10,11,37) Bailey & Doherty, unpublished results).

**Figure 1.**
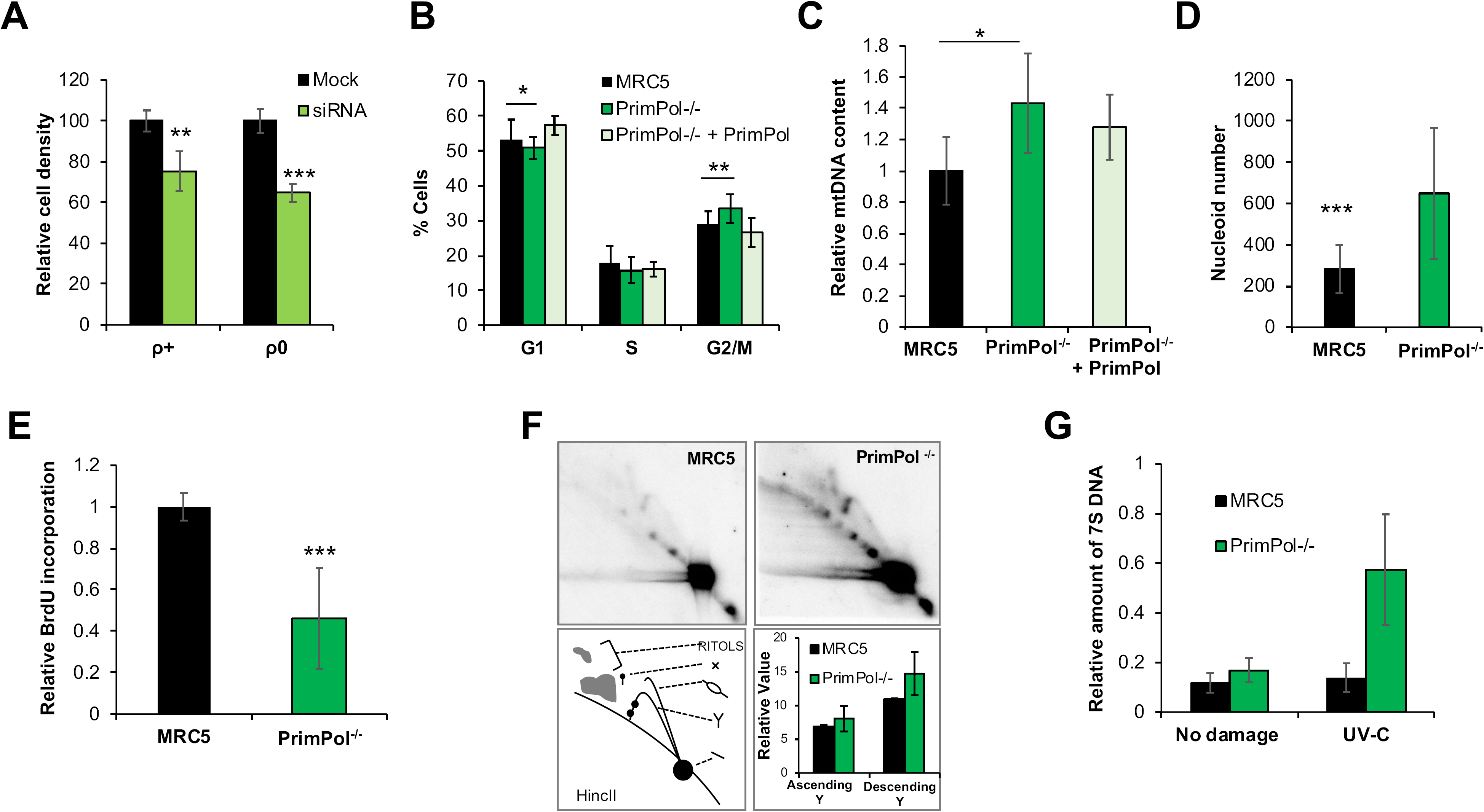
PrimPol knockout cells proliferate normally, have increased mtDNA content but exhibit decreased replication rates. HOS cells treated with PrimPol siRNA or scrambled control were monitored using an IncuCyte^™^ microscope for 80 hrs with cells imaged every 3 hrs and cell density calculated (Fig. S1), (**A**) shows mean cell density at 60 hrs after RNAi from n=3 experiments with standard deviation shown with error bars. (**B**) Cell cycle populations were measured in asynchronously growing MRC5 and PrimPol^-/-^ cells or PrimPol^-/-^ cells complemented with GFP tagged PrimPol after PI staining and flow cytometry. mtDNA copy number was analysed by Taqman probe qPCR in human MRC5 cells (**C**) with and without PrimPol. The mitochondrial network and mtDNA nucleoid were analysed by co-staining cells with Mitotracker (red), anti-DNA (green) and Dapi (blue) and Image J was used to analyse the number of mtDNA nucleoids per cell (**D**). mtDNA replication was measured using BrdU incorporation, nuclear DNA replication was blocked using aphidicolin to allow mtDNA-specific analysis and cells were allowed to accumulate BrdU for 24 hrs. DNA was extracted, dot blotted and BrdU content quantified by anti-BrdU staining, this was then compared to total DNA content by the use of a mtDNA-specific radio-labelled probe, n=3 independent repeats performed in triplicate (**E**). mtDNA replication intermediates were viewed directly by 2D-AGE, total DNA was digested by HinCII and separated in two dimensions. DNA was Southern blotted and probed with an O_H_ specific radio-labelled probe. Images were quantified using AIDA (**F**). 7S DNA copy number was measured in undamaged and UV-C treated cells using southern blotting of whole cell DNA after PvuII digestion and probed with a 7S DNA and nuclear DNA specific radio-labelled probe, and quantified using AIDA software (**G**). Charts represent the mean of n≥3 independent experiments with error bars showing standard deviation, significance was measured using a students t-test, * P≤0.05, ** P≤0.01, *** P≤0.001.

However, the phenotypes of human cell lines transiently depleted of PrimPol significantly contrast with those observed in avian (DT40) PrimPol^-/-^ cells that exhibited increased UV sensitivity, fork slowing and growth arrest after damage (18,19). In order to analyse the roles of PrimPol in mammalian cells in more detail, a human cell line lacking the protein was generated to avoid the variation in PrimPol levels associated with RNAi methods. An engineered gene targeting nuclease (Zfn) (38) was used to generate a DSB within the beginning of the second exon of the *PrimPol* gene, which when repaired inaccurately results in the loss of functional PrimPol. This method allowed us to generate an MRC5 fibroblast cell line lacking PrimPol when analysed by western blotting with a PrimPol-specific antibody (Fig. S2A,B). Reverse transcription PCR (RT-PCR) was also used to analyse transcripts generated by the disrupted gene, confirming that no full-length functional transcripts were observed and sequencing showed significant deletions in both alleles (Fig. S2C). One clone carried a deletion from amino acid 192, with the rest of the gene reading out of frame with a further downstream stop codon, resulting in any potential protein being truncated at residue 239. The second allele carried a deletion of amino acids 182-282 (Fig. S2D). Both deletions result in the loss of at least one key residue of the essential enzyme catalytic triad and would therefore result in large changes in both the protein structure and function. Despite the presence of short gene transcripts, no protein was detected by western blotting using a polyclonal antibody able to recognise all regions of the protein, suggesting that these transcripts are not translated or that any resulting polypeptides are targeted for degradation (16,18)(Fig. S2A,B).

Notably, the resulting PrimPol^-/-^ cells initially showed no obvious signs of distress or growth impediment due to the loss of this enzyme, likely having adapted to the loss of the protein in the time taken (∼ 15 days) to expand the population. Although no differences in cell growth rates were observed, (Fig. S2E), we identified a small but consistent change in cell cycle populations (Fig. 1B), suggesting that there may be underlying consequences due to the loss of PrimPol. We observed a significant increase in the number of cells in G2 / M phase of the cell cycle, with a concomitant loss of those in G1 and S phase. This defect could be complemented by the expression of GFP tagged-PrimPol in PrimPol^-/-^ cells (Fig. 1B). This is similar, but less severe, than the lethal affects observed upon deletion of a PrimPol protein from trypanosomes where cells arrested in G2/M (14).

### PrimPol^-/-^ cells have a defect in mitochondrial DNA proliferation

Previous work on PrimPol^-/-^ mouse embryonic fibroblast (MEFs) highlighted a role for PrimPol in the reinitiation of mitochondrial DNA replication forks after damage (24), therefore we next looked for defects in mitochondrial metabolism and mtDNA maintenance in our human cell line. Mitochondria contain multiple copies of DNA and numbers can vary greatly between tissues and individuals (39). We observed a significant increase in mtDNA copy number (∼ 55 % > than in wild type (WT) cells), in cells lacking PrimPol, which could be complemented by the addition of GFP-tagged PrimPol (Fig.1C). This change was consistent with similar increases observed in a number of different human cell lines treated with PrimPol siRNA (Fig. S3A). In addition, we also found a similar increase (∼ 51%) in mtDNA copy number in DT40 PrimPol^-/-^ cells, which could be complemented by the addition of the WT protein (Fig. S3B), whilst similar increases have also been reported in MEFs (24). Upon staining these cells with an anti-DNA antibody, we found a clear increase in mtDNA nucleoid number but no increase in nucleoid size (Fig. 1D, S3C). Thus, the increase in mtDNA copy number is accommodated by an increase in distinct nucleoids rather than an increase in DNA copies within each nucleoid. Despite the large increase in mtDNA, little change in the size or distribution of the mitochondrial network was observed after cells were stained with Mitotracker, a mitochondrial-specific dye (Fig. S3D). In addition, we found only a minor decrease in mitochondrial membrane potential in PrimPol^-/-^ cells compared with WT MRC5 cells, suggesting mitochondrial function is not unduly impeded after the loss of PrimPol (Fig. S3E).

As PrimPol’s absence causes an increase in mtDNA, we next looked for any changes in mtDNA replication in human cells. To examine this, aphidicolin was used to inhibit nuclear DNA replication and cells were incubated with BrdU for 24 hrs. After DNA extraction and dot blotting, DNA was probed with an anti-BrdU antibody to identify levels of BrdU incorporated over the given time. Total mtDNA was then labelled with a radiolabelled mtDNA probe to account for any loading differences, which may be inherent due to differences in mtDNA copy in relation to total cellular DNA. Strikingly, despite a higher mtDNA copy number, we found a significant decrease in BrdU incorporation as a proportion of total DNA signal in PrimPol^-/-^ cells, demonstrating a large decrease in replication speed or abundance (Fig. 1E). Notably, no gross changes in mtDNA replication intermediates were detected between the two cell lines, as analysed by two-dimensional (2D) agarose gel electrophoresis (Fig. 1F). However, we did identify an increase in the descending arm of the Y-arc, which may be indicative of a slowing in the completion of replication and an accumulation of replication intermediates at later stages of replication. This may indicate a failure to fully complete replication as earlier stages of replication progress unimpeded, as indicated by little differences in the ascending Y-arc. However, replication may stall at later stages where barriers must be bypassed and errors must be repaired, similar to mtDNA replication changes identified in PrimPol^-/-^ MEFS (24). A proportion of mtDNA have a characteristic D-loop, which is a triple-stranded region in the non-coding region containing the origins of replication and transcription. The D-loop is formed by the addition of a short complementary third strand called 7S DNA (40). We identified a small increase in 7S DNA in undamaged PrimPol^-/-^ cells, compared to WT MRC5. However, when cells were treated with UV-C damage, 7S DNA was increased (∼ 3 fold) in PrimPol^-/-^ cells compared with very little change in WT cells (Fig. 1G). This may suggest changes in the mode of replication or initiation as this region contains the origins of replication and transcription, similar to a previous study on PrimPol^-/-^ MEFs that showed replication changes in this region (24).

It was previously reported that, after UV-C damage, mtDNA replication mechanisms switch from a “slow” unidirectional RITOLS mechanism to a “faster” strand coupled mechanism, which would require initiation outside of the D-loop by a poorly understood mechanism (41). Thus, PrimPol may play an important role in allowing replication to be maintained under different conditions (reviewed in (23)). Little is known about the exact role of 7S DNA in mtDNA replication and maintenance but many hypotheses have been suggested, including having a role in replication initiation, serving as a recruitment site for nucleoid proteins or potentially acting as a sink for nucleotides throughout the cell cycle (40,42-44). The increase in 7S DNA after UV-C damage in the absence of PrimPol suggests that one or all of these functions are required at an increased rate. It is feasible that, after damage, all mtDNA molecules may require the recruitment of damage repair proteins to ensure no lesions are left unrepaired or that replication may be delayed after damage in the absence of PrimPol, leading to an increase in mtDNA molecules maintaining their D-loop for longer. In addition, it is also possible that in the absence of PrimPol replication mechanisms required in the presence of Pol γ-stalling lesions are altered, affecting the requirement for D-loop dependent replication initiation. These changes in mtDNA replication and copy number suggest that mtDNA synthesis may be upregulated to overcome defects in electron transport chain (ETC) activity by allowing more copies of mitochondria-encoded proteins to be generated due to a decreased function or increased mutational load in the available copies. In addition, mtDNA replication is a slow process and it is thought that transcription and replication may be unable to occur simultaneously on the same molecule. Thus, any decrease in replication completion rate may have dramatic effects on the viability and functionality of the organelle (23,24).

### PrimPol^-/-^ cells show nuclear replication defects in the presence and absence of DNA damage

Given that PrimPol is also a nuclear protein, we next examined its role in nuclear replication. Using DNA fibre analysis, we measured replication fork rates in WT and PrimPol^-/-^ cells in both the absence and presence of DNA damage. As observed previously in PrimPol^-/-^ DT40 cells, we found that PrimPol-deficient human cells showed a small decrease in replication fork speeds (19) (Fig. 2A). However, this reduction was much less severe than that observed in the mitochondria, likely due to the availability of additional damage tolerance pathways in nuclear DNA replication. In addition, we also observed an increase in sister chromatid exchanges (SCEs) in PrimPol^-/-^ cells, suggesting an increased usage of homologous recombination (Fig. 2B). In combination with the decreased fork speeds, this suggests that PrimPol may play a role in allowing stalled replication to continue, thus preventing the need for alternative restart mechanisms, such as HR. To examine this in more detail, we analysed DNA replication by the induction of exogenously-induced fork stalling lesions by treating cells with UV-C damage. When a high dose of UV was given, between the two labels in the DNA fibre assay, we observed a decrease in the length of the track from the second label, as indicated by an increase in the ratio between the two labels. This increase was more pronounced in PrimPol^-/-^ cells, compared with MRC5 WT cells, and this defect could be complemented by the over-expression of GFP-tagged PrimPol, confirming an increase in fork stalling in the absence of PrimPol (Fig. 2C). This clearly suggests that in the presence of PrimPol forks continue to replicate after damage, whilst in its absence there is a much larger proportion of stalled forks and these must wait for alternative pathways (e.g. TLS) to allow them to be restarted. Together, these data suggest that PrimPol may work directly at the fork, repriming close to the stalling lesion or allowing direct continuation of replication shortly after fork stalling (17,45). As with the other phenotypes compared between the two cell lines, these differences are less significant in human compared to chicken cells, likely due to the increased S-phase population and proliferation rate of avian cells, which duplicate in ∼8 hrs as compared to ∼24 hrs in an average human cells (10,19).

**Figure 2.**
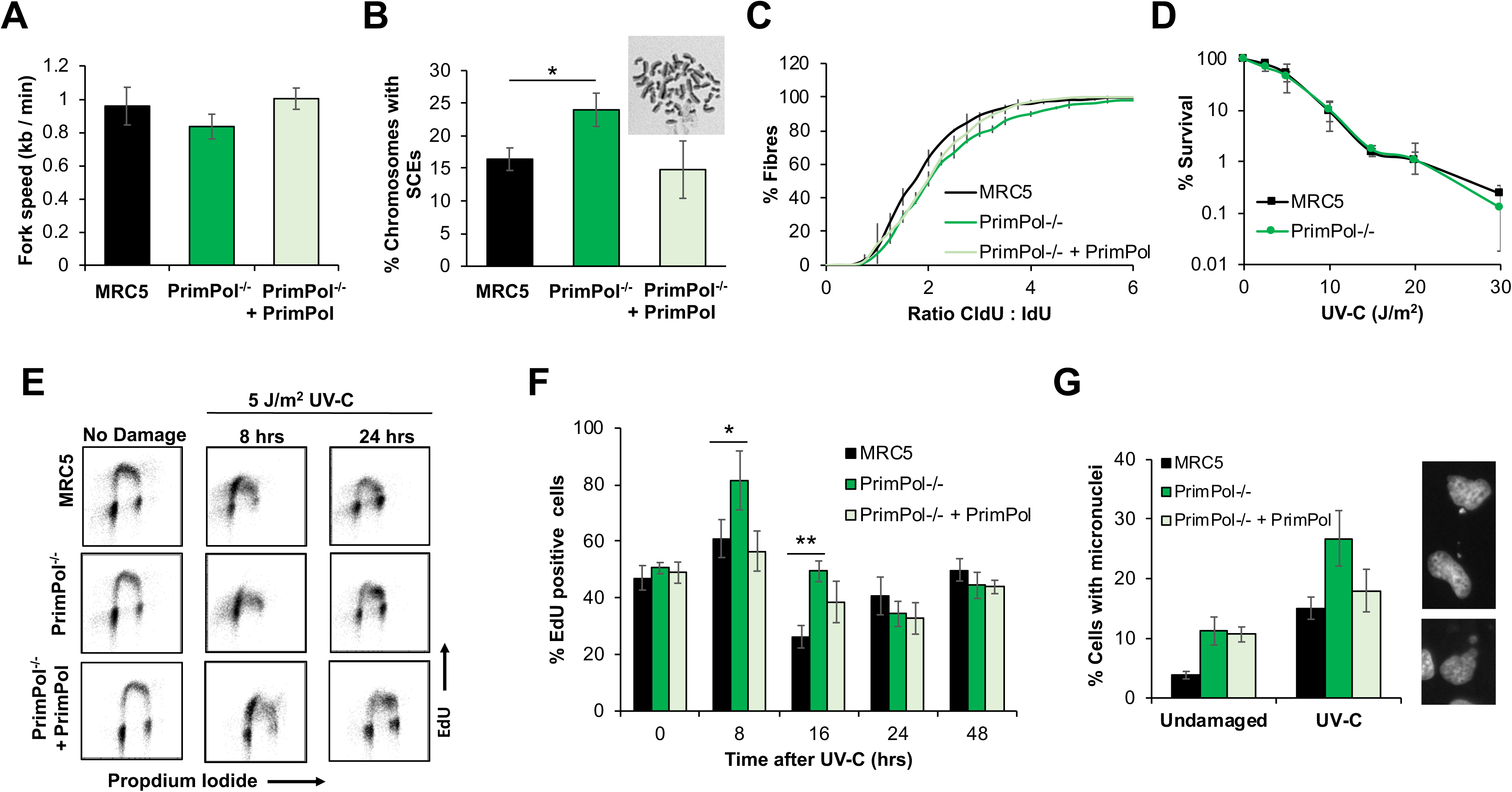
PrimPol^-/-^ cells have decreased replication speeds, increased damage recovery times and micronuclei. (**A**) Replication fork speeds were measured in WT and PrimPol^-/-^ cells or PrimPol^-/-^ complemented with GFP-PrimPol by labelling forks with CldU, followed by IdU and measuring individual fibres after spreading. 300 fibres were measured over 3 independent experiments. (**B**) Percentage of chromosomes carrying one or more sister chromatid exchanges (SCEs) were counted. Data represents 3 independent experiments, where 200+ chromosomes were analysed per slide. Error bars show standard deviation and a representative image of WT cells is shown above. (**C**) This figure shows the accumulated percentage of ratios of the two labels, where a pulse of 20 J/m^2^ UV-C was given in between the two labels. Data is shown as the average across three independent experiments with error bars representing standard deviation. (**D**) Colony survival assays were used to compare the sensitivity of cells to UV-C damage, chart represents n = ≥ 3 independent repeats with error bars showing standard deviation. WT and PrimPol^-/-^ cells or PrimPol^-/-^ complemented with GFP-PrimPol were labelled with EdU at increasing time points after 5 J/m^2^ UV-C damage. After being fixed and dual labelled with propidium iodide, cells were analysed by flow cytometry (**E**). (**F**) shows the percentage of EdU positive cells in early S-phase at increasing times after damage n ≥ 3, error bars show standard deviation. Cells were stained with DAPI 72 hrs after 0 or 5 J/m^2^ UV-C damage and number of cells with 1 or more micronucleus were counted as a percentage of the total cell population, charts show the mean of 3 or more independent experiments with standard deviation shown by error bars. Representative images are shown on the right (**G**).

### Recovery rate, but not survival, is decreased in PrimPol^-/-^ cells after UV damage

Next, we compared the sensitivity of PrimPol^-/-^ and WT MRC5 cells to a range of genotoxic agents. Surprisingly, PrimPol^-/-^ cells exhibited no significant increase in sensitivity to a variety of DNA damaging agents, compared with WT cells (Fig. 2D and S4A-D). This is in stark contrast to PrimPol^-/-^ DT40 cells, which display increased damage sensitivity to many genotoxins, e.g. UV, MMS, 4NQO and cisplatin (18,19). Despite not exhibiting sensitivity, PrimPol^-/-^ cells were clearly impeded to some degree after UV damage as shown by an increase in fork stalling (Fig. 2C), suggesting repair and recovery is slowed in the absence of PrimPol. To examine damage recovery in more detail, EdU labelling and flow cytometry was used to follow cells after UV-C damage. Cells were treated with 5 J/m^2^ UV-C and allowed to recover for up to 48 hrs, at which time all replicating cells were labelled with EdU. DNA was then counter stained with propidium iodide and cell populations analysed by flow cytometry (Fig. 2E, S4E). After UV-C damage, cells accumulated in S-phase as the cell cycle is stalled at the G2 check point to allow repair of damaged DNA and completion of replication. Notably, this stalled population is higher in PrimPol^-/-^ cells than WT counterparts and takes longer to be resolved (Fig. 2F, S4E). At 8 hrs after damage, cells are largely stalled in early S-phase with a slightly larger population and more severe shift towards early S-phase observed in PrimPol^-/-^ cells. Over time, cells progress through S-phase with MRC5 cells returning to a normal cell cycle profile by ∼24 hrs after damage, whilst in PrimPol^-/-^ cells a population of cells is still accumulated in late S-phase at this time, suggesting a delay in recovery / repair in these cells.

While examining PrimPol^-/-^ cells after UV-C damage, we also observed an increase in cells that contained at least one micronuclei (Fig. 2G). Thus, whilst cells are able to recover from UV-C damage, albeit in a delayed manner, they do contain signs that errors or incomplete replication may be a consequence of the loss of PrimPol. Notably, we also observed that micronuclei numbers were increased in PrimPol^-/-^ cells, compared with WT cells, even in the absence of damage. Despite micronuclei being higher in unperturbed PrimPol^-/-^ cells, the further increase observed after damage was still higher in the absence of PrimPol (ratio of damaged / undamaged, 16 for PrimPol^-/-^ versus 11 for MRC5). This is similar to previous observations in DT40 PrimPol^-/-^ cells, where PrimPol was found to be also important for DNA maintenance and replication, even in unperturbed cells (19). These data are suggestive of a delayed recovery from UV-C in PrimPol^-/-^ cells. However, although repair is delayed and damage accumulates, this is not sufficient to cause any major decrease in survival after damage.

### Mutation frequencies are increased in PrimPol^-/-^ cells after DNA damage

To investigate the genetic effects of accumulation of damage in more detail, we used the hypoxanthine phosphorybosyl transferase (HPRT) assay to compare the occurrence of mutations in WT and PrimPol^-/-^ cells after chronic 4NQO damage. Cells were treated with 40 ng/ml 4NQO for 5 days or 10 J/m^2^ UV-C and allowed to grow for a further 6-7 days to allow for activation of any accumulated mutations. Cells were then plated in media with 6-thioguanine, where only those which had gained a mutation or deletion causing inactivation of the *HPRT* gene were able to survive. For all cell lines tested, we found few inactivating mutations in the population in the absence of damage, with mutation frequencies below 4.5 mutants per 10^6^ cells in all cases (data not shown). However, an increase in mutation frequency was observed after damage and, strikingly, levels of drug resistant clones were found to be consistently higher in PrimPol^-/-^ cells, compared to WT cells, suggesting an increase in mutation frequency in the absence of PrimPol (Fig. 3A). This could be complemented by expressing GFP-tagged PrimPol in PrimPol^-/-^ cells. To examine the resulting mutations in more detail, cDNA of the *HPRT* gene from at least 40 clones of each cell line was amplified and sequenced to identify the inactivating mutations. We observed a shift in the type of mutations observed, with an increase in PrimPol^-/-^ cells that had a clear loss of one or more exons from the cDNA (Fig. 3B, C S4F). These changes may be caused by mutations affecting intron / exon splice sites. However, this may also suggest that in the absence of PrimPol, more complex mechanisms than simply bypassing the lesion may be needed, such as HR. Alternatively, the incidence of DSB occurrence may be higher at these sites in the absence of PrimPol due to a delay in restart or repair and thus the break may then be repaired by NHEJ, potentially leading to a loss of genetic information (46,47). Thus, we hypothesise that PrimPol helps with the timely bypass of such lesions, preventing mutation and the loss of vital genetic information. Indeed, it has been shown that in cells depleted of one or more TLS polymerase, the number of non-TLS events, such as big deletions and insertions, increases after UV-C damage (48). However, this assay only identifies mutations that cause a loss of function of the *HPRT* gene so caution must be used in its interpretation as it is likely that a number of synonymous mutations and those which do not cause a loss of function are not identified. Strikingly, when we analysed point mutations found in the *HPRT* gene from each cell line, we found that WT cells predominantly generated transitions, while PrimPol-deficient cells showed an increased accumulation of transversion mutations and this change between the two cell lines was significant when tested using a two way ANOVA P=0.0086 (Fig. 3D, S4F). Interestingly, studies on XPA cells showed that the proportion of transversion mutations are lower after UV-C damage than in undamaged cells suggesting the repair mechanism used avoids these changes (49). Previous work has also shown that a change in the mutation spectrum occurs upon deletion of one or more TLS polymerase, indicating that a change in the lesion bypass mechanism occurs to compensate for the loss of specific damage tolerance pathways (7,48,50,51). The change in mutation signature observed here suggests that a different bypass mechanism is being used in the absence of PrimPol, indicating that PrimPol may play an important role in determining the choice / fidelity of lesion bypass during replication stalling. Thus, it is likely that in the absence of PrimPol, which allows the continuation of replication in the presence of stalling lesions / structures, cells may rely more on canonical TLS pathways to replicate across the impediment, albeit in a more mutagenic way in order to allow completion of replication in a timely fashion. However, when PrimPol is available it likely reprimes replication restart downstream of fork-stalling impediments, allowing TLS pathways to fill in the resulting gaps. This study highlights that, despite its highly mutagenic activity *in vitro* (29), PrimPol actually facilitates replication restart within cells, in conjunction with canonical TLS pathways, using a relatively error-free mechanism. In agreement with this, PrimPol has previously been shown to be important in preventing mutations arising at AP sites in MEFs (37).

**Figure 3.**
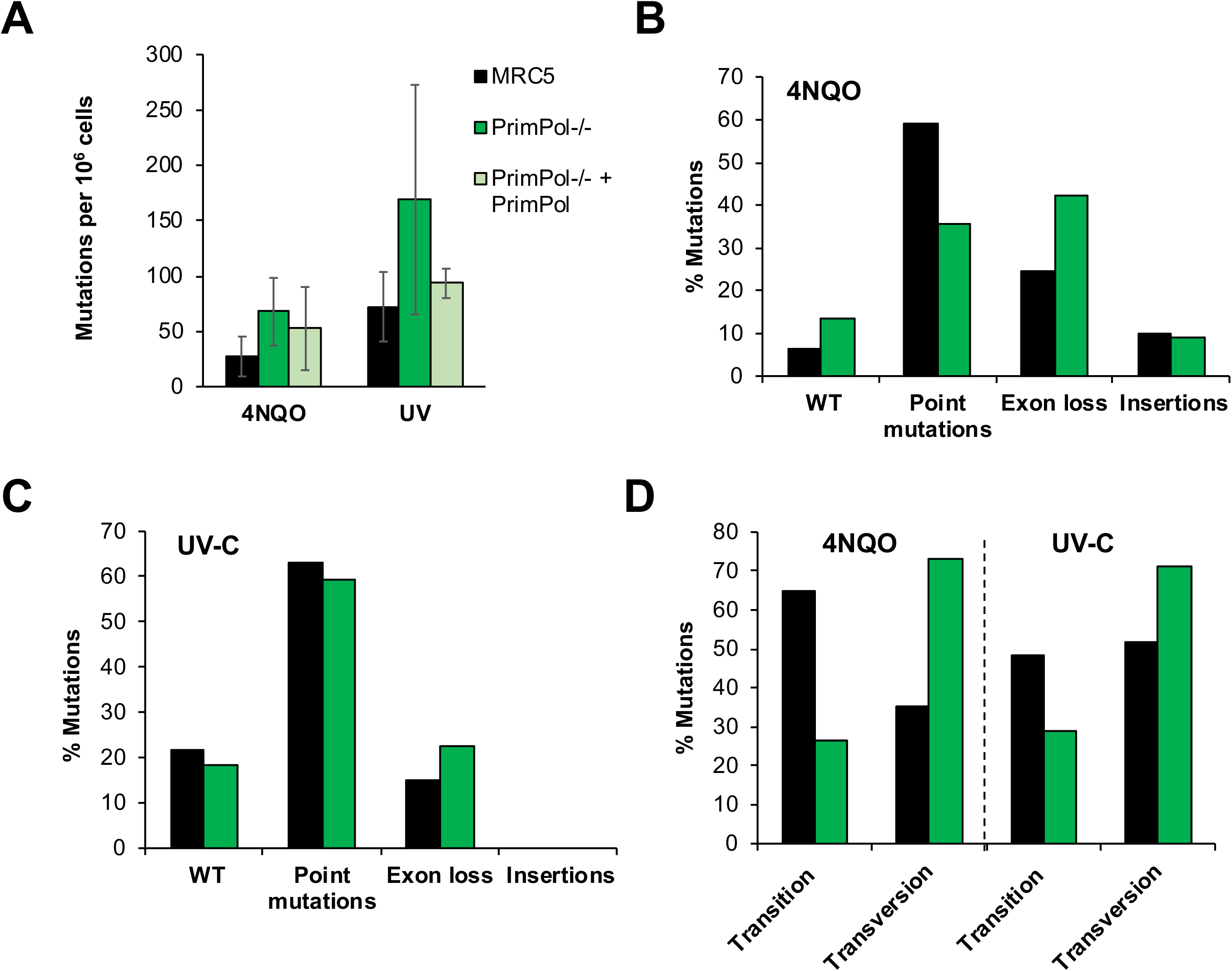
PrimPol^-/-^ cells have increase mutation frequency after damage. Mutation frequencies were calculated after 5 days following 40 ng/ml 4NQO treatment or 10 J/m^2^ UV-C damage, followed by 6-7 days recovery. Cells were plated in the presence of 5 µg/ml 6-Thioguanine to identify clones with a mutation in the *HPRT* gene and the number of colonies were calculated compared to a drug free control plate. Mutation frequency was then calculated as occurrence of a mutation per 10^6^ cells (**A**), bars show the average of ≥ 3 independent experiments and error bars represent standard deviation. RNA was extracted from 6-Thioguanine-resistant clones and the *HPRT* gene cDNA was amplified and sequenced to identify mutations generated after 4NQO treatment (**B**), or UV-C damage (**C**), shown as a percentage of the whole population. Details of the identified clones shown in Fig. S4F. The percentage of point mutations that were transitions or transversions are shown in (**D**) MRC5: PrimPol^-/-^ transitions and transversions P= 0.0086 using two way ANOVA.

### The absence of Pol η highlights the necessity for a PrimPol damage tolerance pathway in eukaryotic cells

In contrast with avian cells, human cells lacking PrimPol exhibit little sensitivity to damaging agents. However, depletion of PrimPol in the Xeroderma pigmentosum variant fibroblast cell line (XPV; XP30RO) increases their sensitivity to UV damage (10). To characterise this phenotype in more detail, we generated a PrimPol^-/-^ deletion in a Pol η-deficient fibroblast cell line (XPV). We first screened for PrimPol loss using western blotting (Fig. S5A), identifying two clones, (XPV PP1 and XPV PP2) lacking any detectable PrimPol protein, compared to a tubulin control. DNA sequencing of these clones identified a clear loss of the ability to produce functional protein in both clones. XPV PP1 contained a missense mutation and a loss of 4 nucleotides (594-597), lead to the remaining portion of the gene being shifted out of frame from amino acid 197 for one allele. The second allele carried a large deletion (140 nt), which begins in the intron before exon 2 and ends at nucleotide 617 in exon 2, causing the loss of an intron exon boundary and much of exon 2. In the second clone, XPV PP2, a 10 nucleotide deletion starting at nucleotide 598 leads the remaining gene to be shifted out of frame, whilst the second allele carries a large insertion at nucleotide 594, which adds a stop codon (Fig. S5B).

Initial analysis revealed little differences in growth rates of cells with the additional loss of Pol η, with doubling times similar to those observed for PrimPol^-/-^ MRC5 cells (Fig. S5C). In addition, XP PP cells also showed a small increase in G2 populations when analysed by flow cytometry using propidium iodide, which could be complemented by the addition of GFP-tagged PrimPol (Fig. 4A). However, unlike in MRC5 cells, loss of PrimPol in XPV cells resulted in a more significant loss of G1 cells and an increase in cells in S-phase, suggesting that replication may be further delayed in unperturbed cells lacking both PrimPol and Pol η.

**Figure 4.**
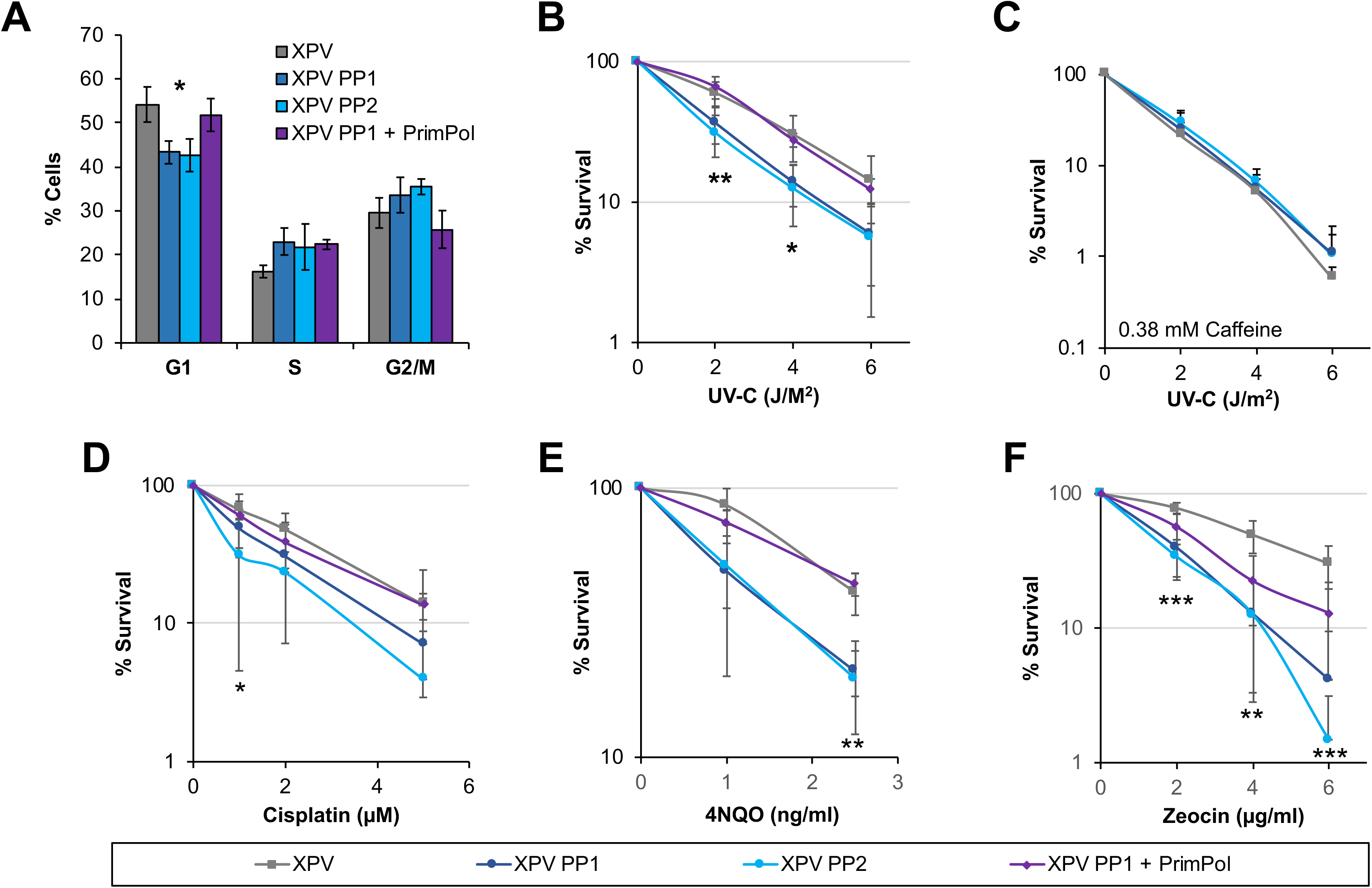
Loss of both PrimPol and Pol η leads to greater damage sensitivity. Cell cycle population changes were studied in WT (XPV) and PrimPol^-/-^ XP30RO cells (XPV PP1 & XPV PP2) and those complemented with GFP PrimPol (XPV PP1 + PrimPol) by flow cytometry analysis of propidium iodide labelled cells (**A**). Sensitivity of cells to a range of DNA damaging agents was measured using colony survival assays (B-F). Cells were treated with increasing doses of UV-in the absence or presence of 0.38 mM caffeine in the media (**B** and **C**), or increasing dose of cisplatin for 5 hrs after which the cells were washed free of the drug (**D**) or cells were maintained in media containing increasing doses of 4NQO or zeocin (**E** and **F**) for 10 days to form colonies before being stained with methylene blue and counted. Charts represent the mean of n≥3 independent experiments with error bars showing standard deviation, significance was measured using a students t-test, * P≤0.05, ** P≤0.01, *** P≤0.001.

As Pol η is a TLS polymerase important for the bypass of CPDs and PrimPol has the ability to bypass 6-4 photoproducts using TLS (10), we next studied the sensitivity of the double-deletion cells to UV-C exposure. Unlike WT or PrimPol^-/-^ MRC5 cells, XPV PP cells showed a significantly greater UV-C sensitivity than XPV cells alone (Fig.4C), similar to previously observed using RNAi depletion (10). XPV cells have been reported to display increased sensitivity to UV-C but this is dependent on the addition of low doses of caffeine (52), thus UV-C colony survival assays were repeated in the presence of 0.38 mM caffeine. Strikingly, only a minor increase in sensitivity was observed in XPV PP cells but XPV cell survival was significantly reduced to the same level as the double knockout cells, suggesting that the pathway inhibited by low caffeine levels may involve PrimPol (Fig. 4C). Low levels of caffeine treatment after UV-C damage has been shown to cause premature chromatin condensation, along with an increase in the presence of ssDNA after collapse of the replication fork and a decrease in Chk1 phosphorylation (53-55). Here, we observed that loss of PrimPol causes little additional sensitivity, when low levels of caffeine were present in colony survival assays. This suggests that, in the presence of caffeine, PrimPol may be unable to function to maintain replication forks either because its activity is decreased, replication forks may become inaccessible to the protein or it may fail to be recruited in the correct way, possibly due to a decrease in checkpoint activity.

In addition, XPV PP cells also showed sensitivity to other DNA damaging drugs. These included cisplatin and zeocin that induce DNA crosslinks and DSBs, respectively (Fig. 4D, F). In all cases, this sensitivity could be complemented by the addition of GFP-tagged PrimPol. Together, these findings further support the proposal that PrimPol forms part of a complementary damage tolerance pathway that allows the replication machinery to bypass different types of DNA lesions.

In contrast, although XPV cells alone showed higher levels of micronuclei than MRC5 WT cells, no additional increase was observed upon the loss of PrimPol (Fig. S5D). This was also true in the case of UV-C damage, where XPV cells again showed elevated levels of micronuclei but no further increase with the loss of PrimPol. This may be explained by a change in the levels of accumulated damage and / or, as indicated by the observed decreased survival, a failure to appropriately complete replication in time that leads to a significantly increased damage load, ultimately resulting in cell death. Unfortunately, we were unable to analyse the change in mutation spectrum in the XPV PP cells due to an additional X chromosome in the parental cell line (56).

To examine cell recovery after UV-C damage in more detail, cells were labelled with the dNTP analogue EdU and their cell cycle progression followed by flow cytometry. A lower UV-C dose of 2 J/m^2^ was used due to the greater UV-C sensitivity of the XPV cells. As observed in MRC5 cells, recovery was clearly slowed in the absence of PrimPol^-/-^, but this defect was even further prolonged with the additional loss of Pol η (Fig. 5A, quantified in Fig. 5B, S6A,B). Whereas XPV cells had largely completed S-phase at 24 hrs after damage, those also lacking PrimPol were still stalled in early S-phase, suggesting that many cells were unable to overcome fork-impeding UV-C induced lesions and must utilise slower repair and / or bypass mechanisms. This is likely to account for the additional sensitivity observed in survival assays as many of these cells are unable to overcome and successfully repair this damage, likely resulting in cell death.

**Figure 5.**
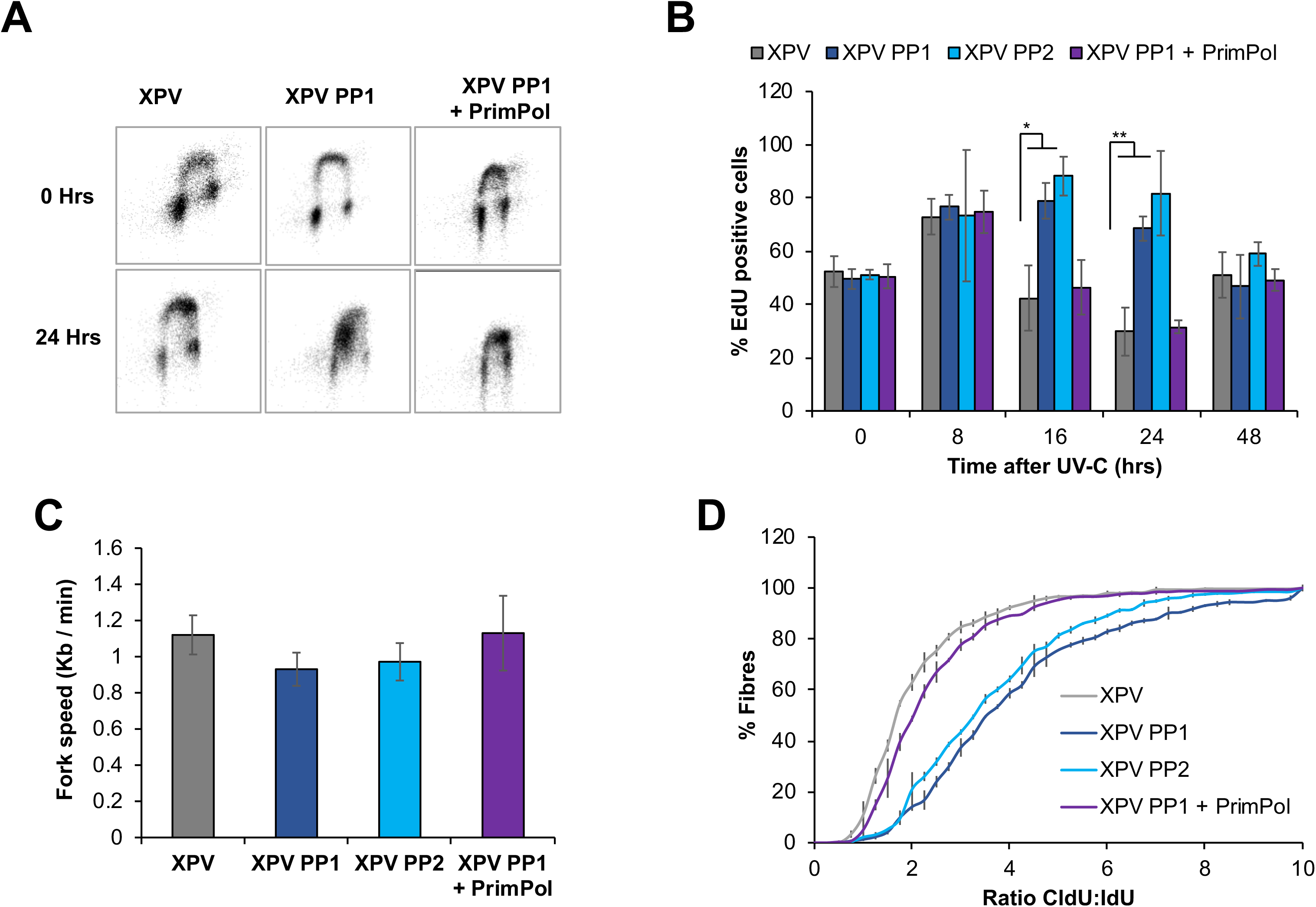
XPV PP cells have an increase in replication defects. Cells were treated with 2 J/m^2^ UV-C and allowed to recover for increasing times before being labelled with EdU. Cell cycle populations were then measured by the addition of propidium iodide, representative images are shown in (**A**) with percentage of EdU cells in early S-phase quantified in (**B**). Replication fork speeds were measured in undamaged cells by labelling with CldU followed by IdU, after spreading and staining with specific antibodies, fibres were measured using OMERO, 300 fibres were measured over 3 independent experiments (**C**). Fork stalling was measured by the addition of a 20 J/m^2^ pulse of UV-C, between the addition of the two labels, and the ratio of the two labels was compared and is shown as the accumulated percentage of the average of each experiment (**D**). Charts represent the mean of n≥3 independent experiments with error bars showing standard deviation, significance was measured using a students t-test, * P≤0.05, ** P≤0.01, *** P≤0.001.

We next looked at the effect of loss of both PrimPol and Pol η on replication fork speeds. No differences in fork speed were observed in XPV cells, compared to WT MRC5 cells, but a clear loss of fork speed was noted with the additional loss of PrimPol, similar to PrimPol^-/-^ single knockout cells (Fig. 5C). PrimPol^-/-^ cells showed a ∼12.8 % decrease in fork speed, compared with WT MRC5 cells, whereas although XPV cells actually had ∼17 % higher replication fork speed compared with MRC5 cells, they also showed a similar decrease in fork speed in cells also lacking PrimPol, with clones 1 and 2 decreased by 19.8 and 13.2 %, respectively. This suggests that PrimPol plays an important role in the maintenance of replication fork progression under normal unperturbed conditions, whereas Pol η may be dispensable under such circumstances, only becoming requisite after the addition of exogenous damage. However, when replication fork progression was measured after UV-C damage, a much more significant affect was identified (Fig. 5D). Notably, XPV cells showed only a small increase in fork stalling after UV-C damage, compared to WT MRC5 cells. However, a significant increase in replication fork stalling was observed when both Pol η and PrimPol were absent. These changes are similar to those observed in survival assays and cell cycle progression, indicating that the cause of the cell cycle delay observed after damage is a failure of replication fork progression after encountering damage on the template strand. By deleting PrimPol in the absence of Pol η, we have uncovered an overlapping, but non-epistatic relationship between these two complementary damage tolerance pathways, highlighting the importance of PrimPol for maintaining efficient nuclear DNA replication in human cells, even in the absence of exogenous damage.

## Acknowledgements

Thanks to Essen Biosciences for the use of the IncuCyte^™^ microscope. Prof. Alan Lehmann for the XP30RO cells. AJD’s laboratory supported by grants from the Biotechnology and Biological Sciences Research Council (BBSRC: BB/H019723/1 and BB/M008800/1).

We declare that none of the authors have a financial interest related to this work.

---

**Supplementary Figure 1.**
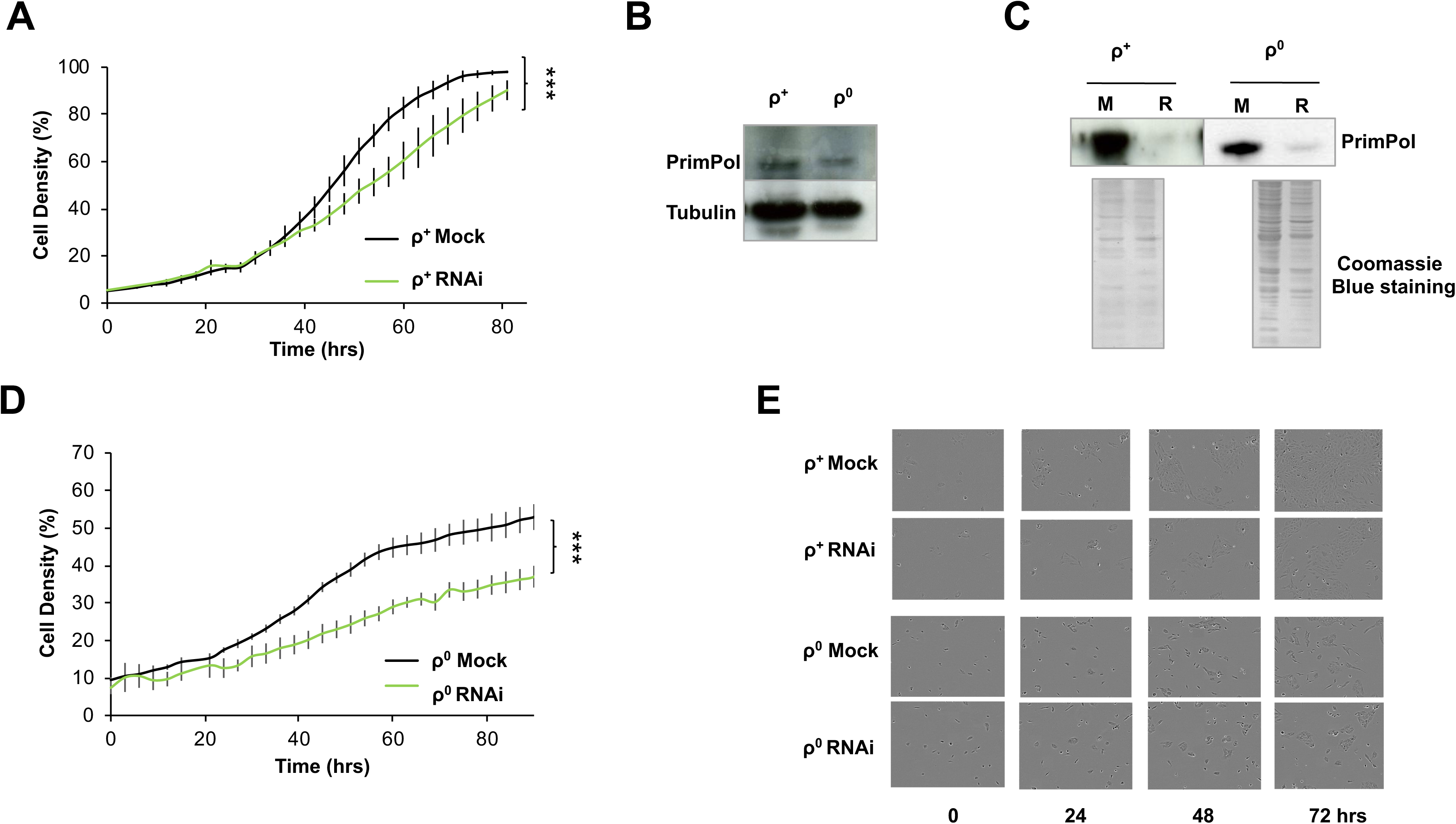
PrimPol RNAi causes a decrease in cell proliferation in both WT and ρ^0^ cells. An example growth curve for 143B cells following transfection with siRNA targeted at *PrimPol* or a scrambled control. Images were taken using an IncuCyte^™^ phase contrast microscope and chart represents an average of two or three individual cell samples grown concurrently with 9 regions analysed per sample at 3 hourly time points error bars show standard deviation across different samples (**A**). PrimPol levels were analysed by western blot of whole cell lysate in wild type (ρ^+^) human osteosarcoma cells and those lacking mtDNA (ρ^0^), quantified in relation to tubulin levels (n=3) (**B**). PrimPol depletion was confirmed 72 hrs after RNAi treatment (R), compared with mock treated cells (M) by western blot of whole cell lysate with comassie to show similar levels of loading (**C**). Example growth curves for ρ^0^ 143B cells after PrimPol RNAi treatment at time 0 taken using an IncuCyte^™^ phase contrast microscope (**D**). Each growth curve represents an average of two or three individual cell samples grown concurrently with 9 regions analysed per sample at 3 hourly timepoints. (**E**) shows representative images taken from the microscope at the specific timepoints.

**Supplementary Figure 2.**
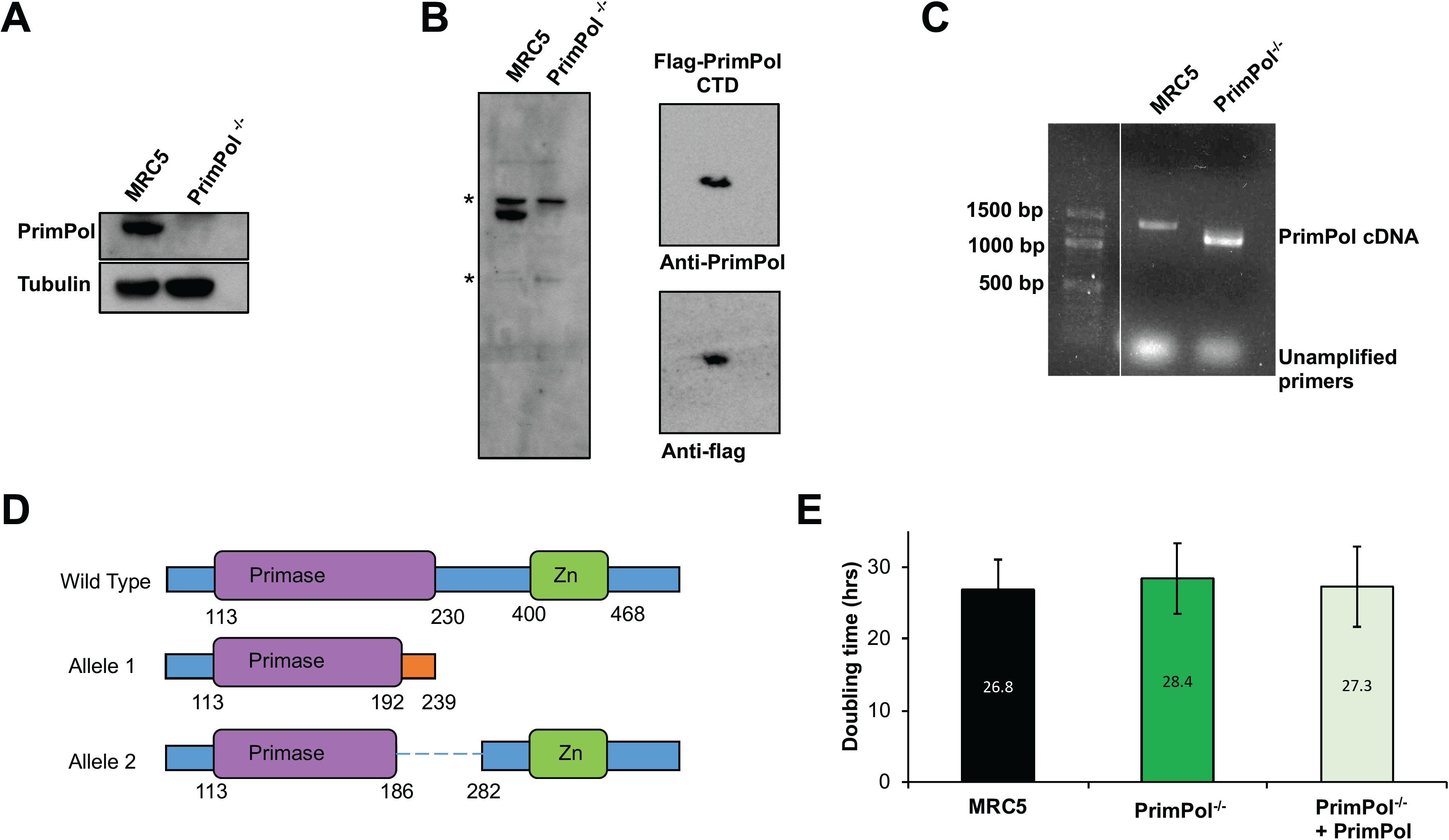
Generation of a PrimPol human knockout cell line. Generation of MRC5 PrimPol^-/-^ cells was confirmed using western blotting of whole cell lysate with a PrimPol specific antibody (**A**) against a tubulin control. (**B**) shows the extended western along with a western showing identification of a c-terminal truncated form of PrimPol (**C**) rtPCR analysis identified only truncated forms of PrimPol mRNA in PrimPol^-/-^ cells. Genomic DNA changes were identified by sequencing and these are depicted in cartoon form, amino acid numbers shown below (**D**). Doubling time was analysed by counting cell numbers over increasing time in both WT and PrimPol^-/-^ cells or those complimented with GFP PrimPol, n≥3 with standard deviation shown as error bars (**E**).

**Supplementary Figure 3.**
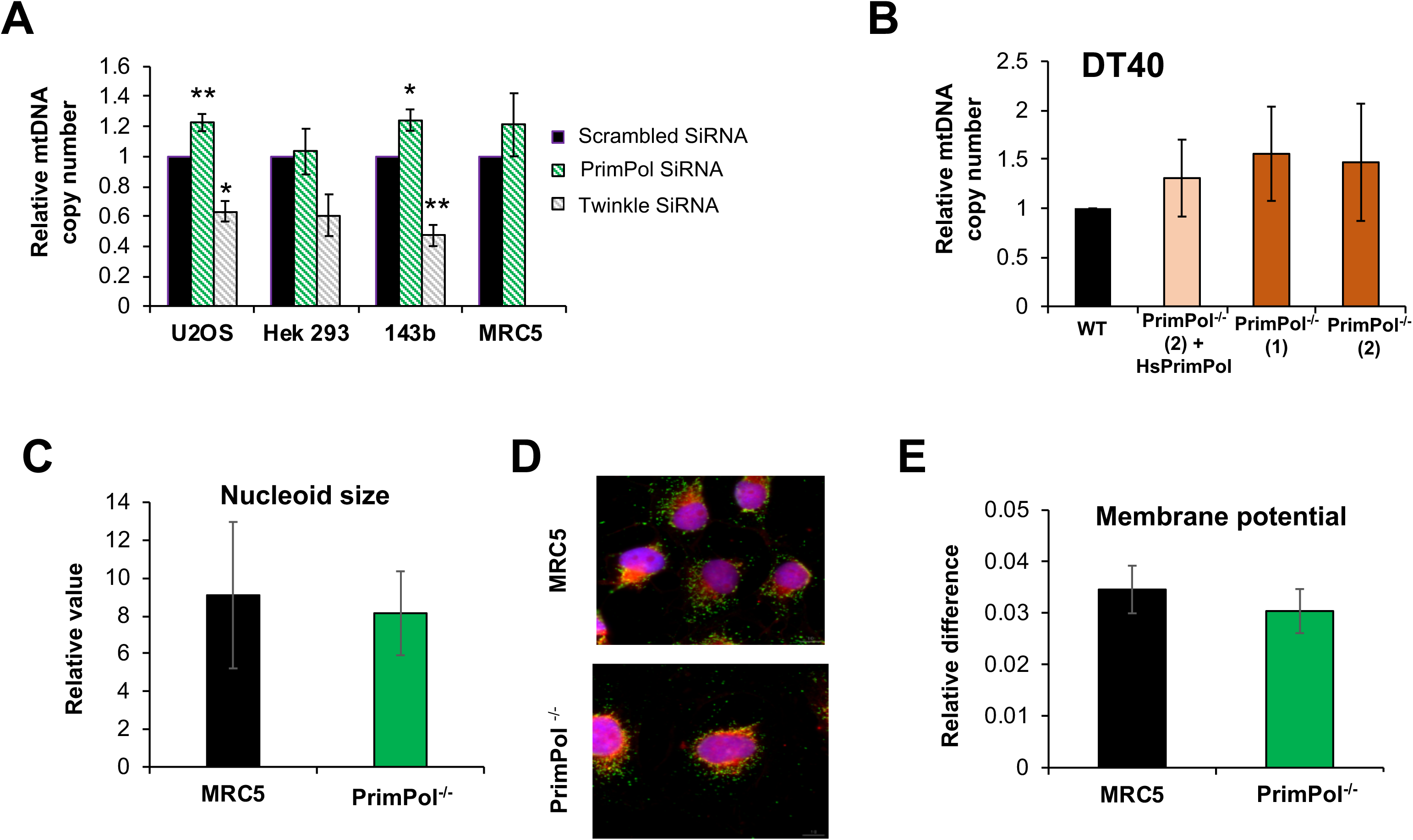
mtDNA copy number is increased in cells lacking PrimPol along with other replication changes. Different cell lines were treated with siRNA targeted to PrimPol or Twinkle or scrambled siRNA and mtDNA copy number was analysed by QPCR in comparison to the scrambled control (**A**), data is the mean of n≥3 (MRC5 n=2) with standard deviation represented by error bars. mtDNA copy number was also compared in WT DT40 chicken cells, 2 PrimPol^-/-^ DT40 clones or those complimented with HsPrimPol (described in (10)) (**B**). PrimPol^-/-^ and MRC5 control cells were used to analyse mtDNA organisation and functionality, nucleoid size was measured using Image J cells stained with Mitotracker and anti-DNA (**C**), representative images shown in (**D**), Mitotracker stained mitochondria, red, mtDNA nucleoids stained with antiDNA antibody green and nucleus DAPI, shown in blue. Mitochondrial membrane potential was measured using Mitotracker by comparing staining with a mass specific, membrane potential independent dye and a membrane dependent dye using flow cytometry (**E**).

**Supplementary Figure 4.**
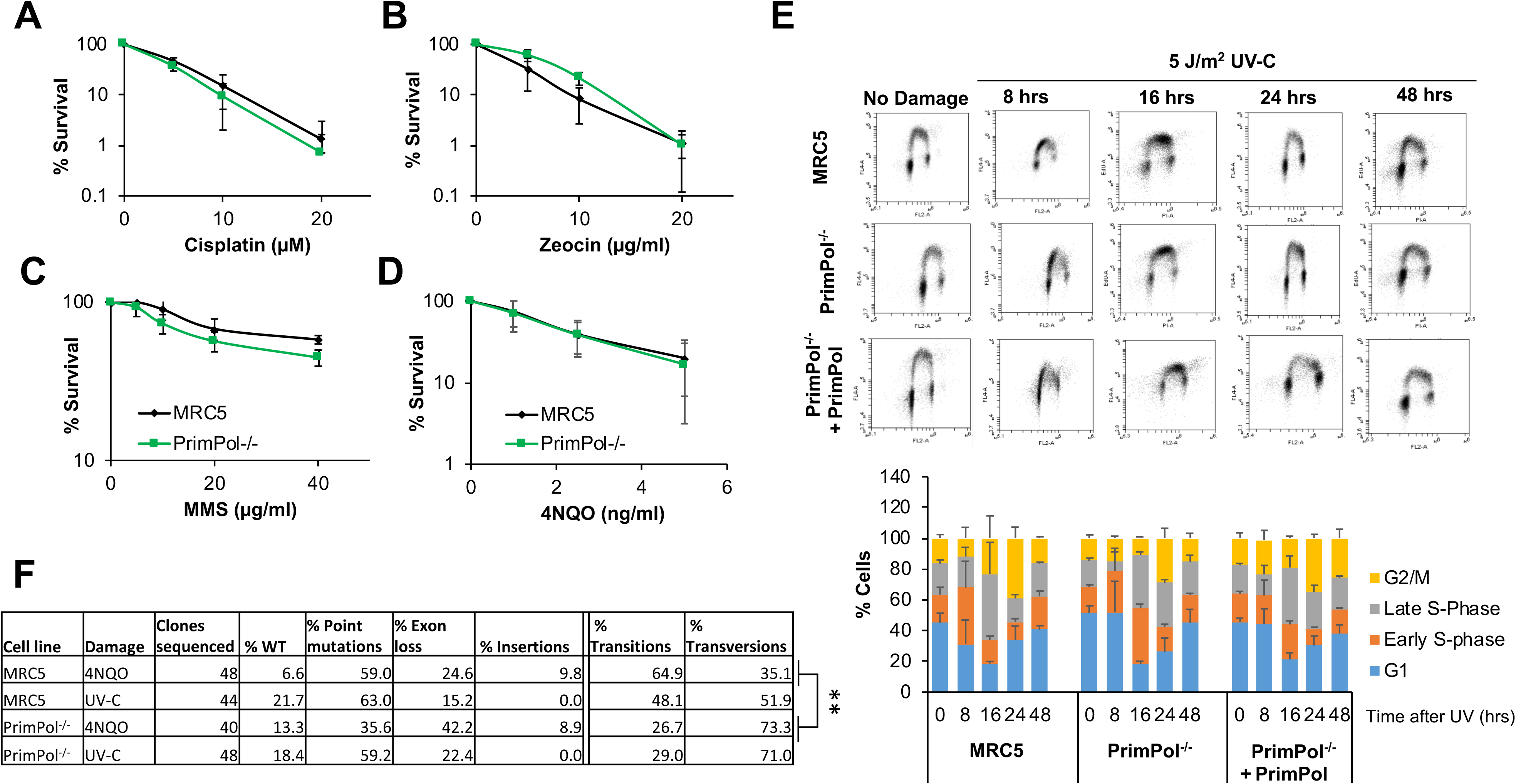
PrimPol^-/-^ cells are not sensitive to DNA damaging agents but have an increased recovery time. PrimPol^-/-^ cells showed no significant increase in sensitivity when treated with a range of DNA damaging agents, Cisplatin, Zeocin, methyl methanosulphate (MMS) and 4NQO. (**A**, **B**, **C**, **D**), charts represent n=3 or more independent experiments with standard deviation shown as error bars. (**E**) Representative images of flow cytometry of EdU and propidium iodide stained cells at increasing time points after 5 J/m^2^ UV-C damage quantified in Figure 2F, full quantification of the cell cycle profiles shown in (**F**) n= 3 with standard deviation shown as error bars. (**G**) shows details of the *HPRT* clones screened for mutations. P= 0.0086 in a two way ANOVA for changes in transitions and transversions between PrimPol^-/-^ and WT MRC5 cells after damage.

**Supplementary Figure 5.**
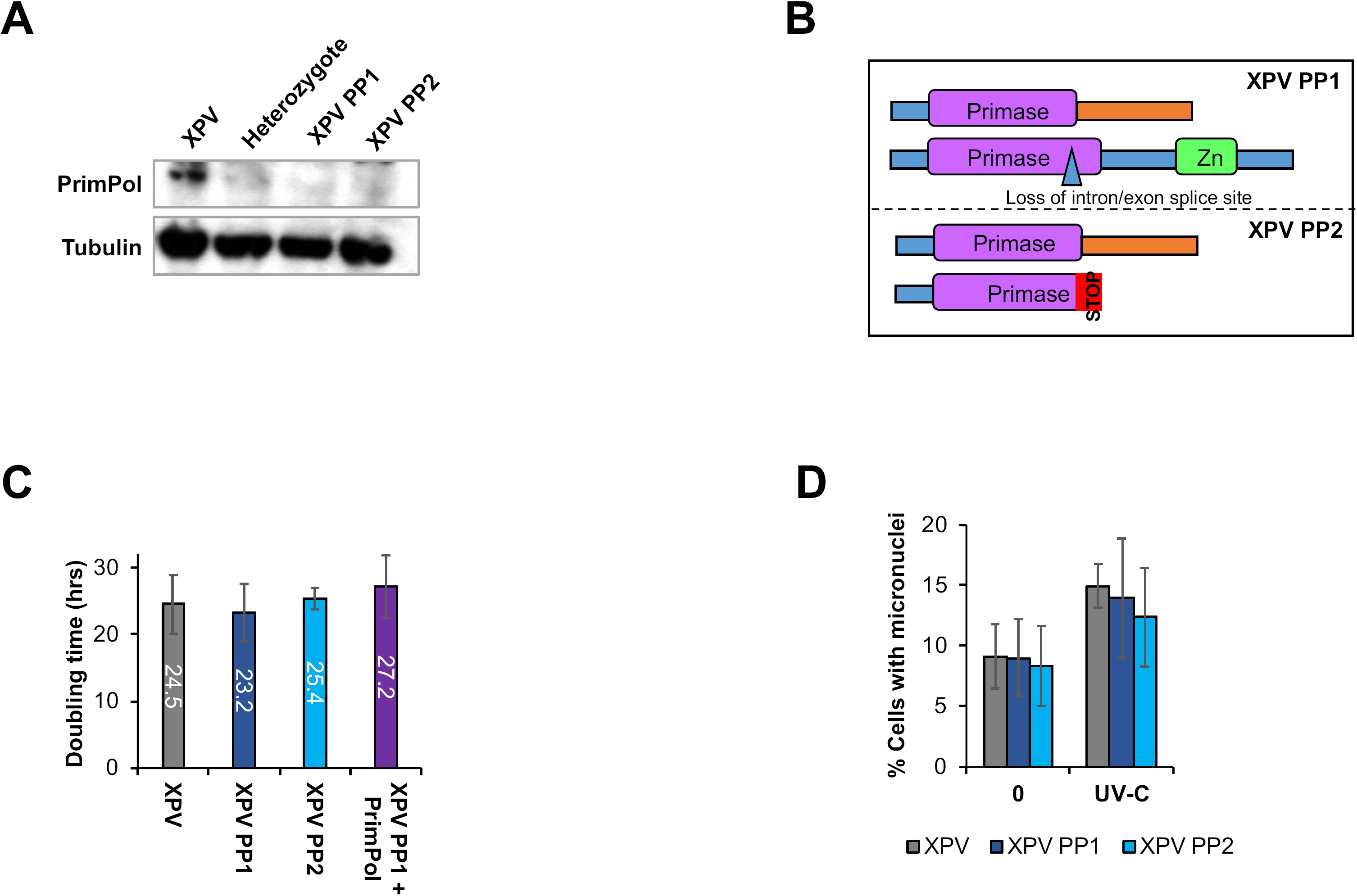
Generation of PrimPol^-/-^ cells in the Pol η^-/-^ XP30RO cell line. Two XP30RO clones lacking PrimPol protein expression were identified by western blot of whole cell lysate with a PrimPol specific antibody along with tubulin control, XP PP1 and XP PP2 (**A**). (**B**) shows the genomic changes observed in the two PrimPol^-/-^ clones in the XP30RO causing the loss of the protein. Growth rates were analysed in WT XPV cells along with the PrimPol^-/-^ clones and those complimented with GFP PrimPol by counting on a hemacytometer (**C**). Cells were stained with DAPI 72 hrs after 0 or 2J/m^2^ UV-C and cells with 1 or more micronuclei were counted as a percentage of the whole population (**D**).

**Supplementary Figure 6.**
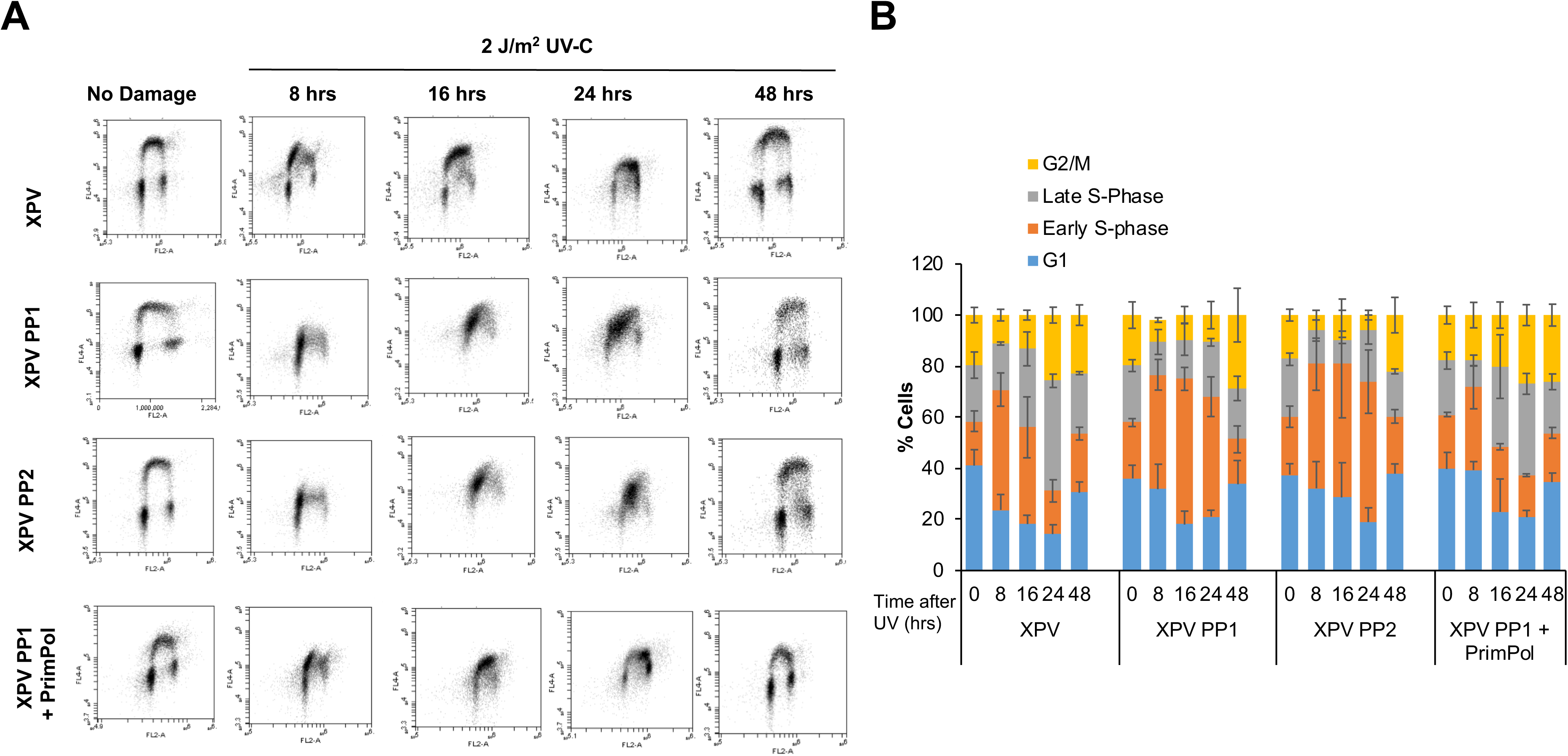
XP PP cells have delayed recovery after UV-C damage. (**A**) Representative images of flow cytometry analysis of cells stained with EdU at increasing time points after 2 J/m^2^ UV-C, quantified in Fig. 5B. Full quantification of the cell cycle profiles shown in (**B**) n ≥ 3 with standard deviation shown as error bars.

